# The Penultimate Step of Proteasomal ATPase Assembly is Mediated by a Nas2-Dependent Switch

**DOI:** 10.1101/2022.10.25.513763

**Authors:** Suganya Sekaran, Soyeon Park

## Abstract

The proteasome holoenzyme is a complex molecular machine that degrades most proteins. In the proteasome holoenzyme, six distinct ATPase subunits (Rpt1 through Rpt6) enable protein degradation by injecting protein substrates into it. Individual Rpt subunits assemble into a heterohexameric “Rpt ring” in a stepwise manner, by binding to their cognate chaperones. Completion of the heterohexameric Rpt ring correlates with release of a specific chaperone, Nas2, however it is unclear whether and how this event may ensure proper Rpt ring assembly. Here, we examined the action of Nas2 by capturing the poorly characterized, penultimate step of heterohexameric Rpt ring assembly. We show that Nas2 uses steric hindrance to block premature progression of the penultimate step into the final step of Rpt ring assembly. Importantly, Nas2 can activate an assembly checkpoint via its steric activity, when the last ATPase subunit, Rpt1, cannot be added in a timely manner. This checkpoint can be relieved via Nas2 release, when it recognizes proper addition of Rpt1 to one side of its cognate Rpt5, and ATP hydrolysis by Rpt4 on the other side of Rpt5, allowing completion of Rpt ring assembly. Our findings reveal dual criteria for Nas2 release, as a mechanism to ensure both the composition and functional competence of a newly assembled proteasomal ATPase, to generate the proteasome holoenzyme.

## INTRODUCTION

The proteasome holoenzyme is responsible for degrading most proteins in the cell. It forms via association of three sub-complexes: the 9-subunit base, 9-subunit lid, and 28-subunit core particle (CP) (1,2). In the proteasome holoenzyme, the base sub-complex recognizes polyubiquitinated proteins, unfolds and translocates them into the barrel-shaped CP, where their degradation occurs (3,4). During this process, the lid cleaves the polyubiquitin chain from the protein substrate, facilitating substrate translocation into the CP. These multi-step events of protein degradation occur through a series of specific conformational changes between the base and lid, and also relative to the CP (5–9). The base coordinates these conformational changes using ATP hydrolysis through its six distinct ATPase subunits (Rpt1 through Rpt6), which are arranged into a ring-shaped complex, forming a key structure of the base (6–10). The heterohexameric ATPase “ring” provides the lateral binding site for the lid, generating a base-lid complex referred to as the regulatory particle (RP) (10,11). In the RP, the heterohexameric ATPase ring docks into the cylindrical end of the CP, mediating RP-CP association to complete proteasome holoenzyme assembly (6–10). RPs can associate with either one or both ends of the CP, generating a singly- or doubly-capped proteasome holoenzyme termed RP_1_-CP or RP_2_-CP (1,2). These features of the base emphasize its importance for both assembly and function of the proteasome holoenzyme.

Base assembly relies on four dedicated chaperone proteins, Rpn14, Nas6, Hsm3 and Nas2, which are conserved between yeast and humans (12–18). Each specific chaperone acts by binding to its cognate Rpt protein: Rpn14-Rpt6, Nas6-Rpt3, Hsm3-Rpt1, Nas2-Rpt5 (12–14,16,17). Three distinct modules then form, with each module containing one or two chaperones: Rpn14-Rpt6-Rpt3-Nas6, Hsm3-Rpt1-Rpt2-Rpn1, and Nas2-Rpt5-Rpt4. Chaperones together facilitate the three modules to associate into a heterohexameric Rpt ring to complete the base complex, and further regulate association of the base with the lid and CP to generate the proteasome holoenzyme. In the current model, binding of each specific chaperone to their cognate Rpt protein sterically hinders the incoming lid or CP, until base assembly is complete, to prevent formation of defective proteasome complexes (15,16,19–23). Upon completion of the proteasome holoenzyme, the chaperones are evicted (15,16,20,24).

Although the base complex *en route* to the proteasome holoenzyme contains 3 chaperones (Rpn14, Nas6 and Hsm3), it excludes a fourth chaperone, Nas2, by releasing Nas2 specifically (25–27). This correlation that Nas2 releases from the assembled base is thought to be a potential checkpoint for base assembly, due to steric conflict between Nas2 and the fully-formed base (25). However, this aspect of base assembly via Nas2 has not been examined experimentally. Contrary to multiple approaches for examining the base-binding chaperones (Rpn14, Nas6 and Hsm3) during the progression of the assembled base into the proteasome holoenzyme (12,16,17,19,20,22,24,28), approaches for examining the Nas2 chaperone have been challenging to establish, due to the rapid progression and the low level of the assembly intermediates, prior to completion of the base. One of the least characterized segments of base assembly is the penultimate step involving Nas2. At this step, the Nas2-bound Rpt5-Rpt4 module is proposed to associate with a rate-limiting Rpn14-Rpt6-Rpt3-Nas6 module, which is considered as a “seed” of base assembly but has not yet been detectable in yeast (17,26). The resulting complex is then thought to incorporate the Hsm3-Rpt1-Rpt2-Rpn1 module, completing base assembly and releasing Nas2 (25,26). Our studies here examine this poorly characterized segment of base assembly to investigate whether and how Nas2 may ensure proper completion of base assembly.

In the present study, we examined Nas2 actions by capturing the penultimate step of base assembly using both a heterologous *E. coli* system and budding yeast, *S. cerevisiae*. We find that Nas2 sterically hinders addition of other subunits until the penultimate base forms. Nas2 allows for completion of the base complex, specifically when ATP hydrolysis by Rpt4 occurs, providing a signal for Nas2 release and proper incorporation of Rpt1. Through this mechanism, Nas2 may ensure both subunit composition and functional competence of the newly assembled base for generating a proteasome holoenzyme.

## RESULTS

### Isolation of the penultimate base complex using a heterologous *E. coli* system

To investigate early-stage base assembly events, which occur too rapidly to capture in yeast, we used a heterologous system in *E. coli* co-expressing 9 base subunits and 4 assembly chaperones (29). In this system, two different affinity tags, one FLAG and six histidines (His_6_), are appended to Rpt1 and Rpt3, respectively. A functional base can be isolated via two consecutive affinitypurifications, His6-Rpt3 followed by FLAG-Rpt1 as a bait, since these two Rpt proteins form two different modules first and later join to complete the base complex (1,2,29). Here, we took advantage of this system and conducted two parallel His6-Rpt3 vs. FLAG-Rpt1 affinity-purifications, to obtain assembly intermediates that are *en route* to the base but are difficult to isolate due to their low level or rapid progression during endogenous base assembly.

Native gel analysis of FLAG-Rpt1 affinity-purified materials exhibited one major complex. This complex is the fully-formed base, as confirmed by a proteomics analysis (Fig. 1, lane 1; see Table S1). All 9 integral base subunits were identified, together with 3 assembly chaperones (Nas6, Rpn14 and Hsm3). The fourth chaperone, Nas2, is absent in the base, consistent with the evidence suggesting that Nas2 releases prior to completion of the base (Fig. 1) (25–27). In addition to the fully-formed base complex, a trace amount of the other Rpt1-containing complex, the Hsm3-Rpt1-Rpt2-Rpn1 module, was detectable by immunoblotting (Fig. 1, lane 1; Fig. S1, lane 1). These results are consistent with the established order of base assembly steps, where Rpt1 exists mainly in 2 complexes: the Hsm3-Rpt1-Rpt2-Rpn1 module and the fully-formed base (12,14,17).

**Figure 1.**
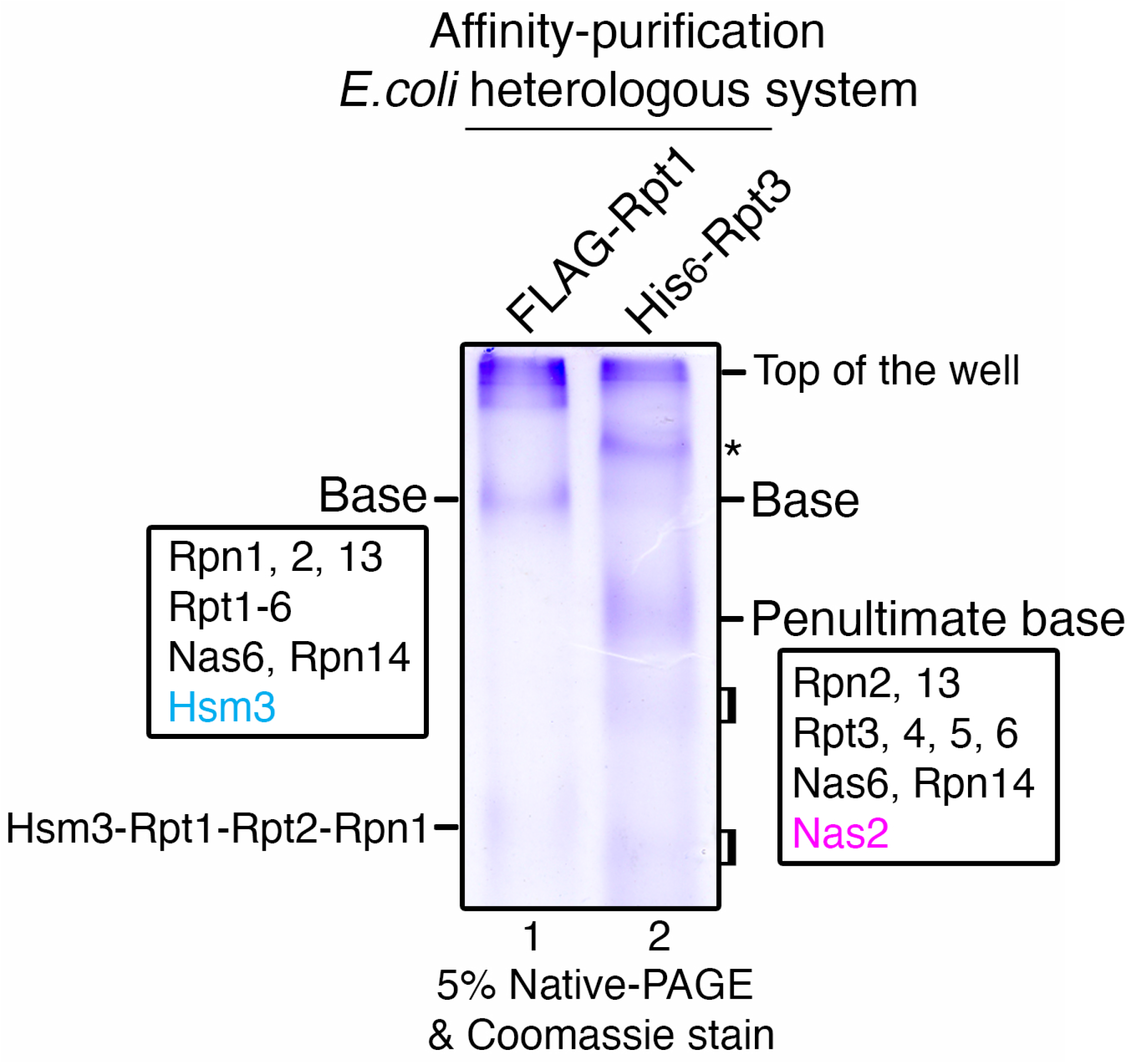
Isolation of the penultimate base complex using the heterologous *E. coli* system. The penultimate base can be affinity-purified using His6-tagged Rpt3 as a bait in the heterologous *E. coli* system, together with the base, which can be isolated using FLAG-tagged Rpt1 (29). Purified proteins (10 μg) were subjected to 5% native-PAGE, followed by Coomassie blue staining. Subunit composition of the base and penultimate base was identified by excising the corresponding bands for a proteomics analysis (Table S1). Two broad, faint bands below the penultimate base (brackets, lane 2) are not reactive with Rpt6, suggesting that they are not likely to be the Rpn14-Rpt6-Rpt3-Nas6 module (Fig. S1, lane 2).

On the other hand, our native gel analysis of His6-Rpt3 affinity-purified material exhibited several bands (Fig. 1, lane 2). The slowest-migrating band was identified as non-specific *E. coli* proteins (Fig. 1, asterisk, lane 2) and the faint band underneath it is the base complex, as seen in FLAG-Rpt1 affinity-purification (Fig. S1, lane 2). Importantly, a more prominent band below the base was identified as a 9-subunit complex, consisting of 2 non-ATPase subunits (Rpn2 and Rpn13), 4 ATPase subunits (Rpt3, 4, 5, 6), and 3 chaperones (Nas6, Rpn14, Nas2) (Fig. 1, lane 2 and Table S1; also see Fig. S1, lane 2). Since this complex contains all base components, except the last module, Hsm3-Rpt1-Rpt2-Rpn1, whose incorporation completes base assembly, we will refer to it as the penultimate base (“pu-base” in short) henceforth.

The penultimate step of base assembly has been poorly characterized, due to its low steadystate level and rapid progression during endogenous proteasome assembly. Now that we can capture the penultimate base complex (Fig. 1, lane 2), we can examine how this step contributes to proper completion of base assembly. In particular, an existing model proposes that this transition into the final step of base assembly relies on the Nas2 chaperone as a potential checkpoint, such that Nas2 may sterically hinder the penultimate base from prematurely progressing into the base complex (25). Upon completion of the base complex, Nas2 may then release (Fig. 1, lane 2 vs. 1). We sought to test the proposed action of Nas2 using the heterologous system, since the penultimate and fully-formed base can be isolated in parallel, through His6-Rpt3 and FLAG-Rpt1 affinity-purification, respectively (Fig. 1).

### Nas2 blocks the premature progression of the penultimate base into the base

If Nas2 provides a checkpoint at the penultimate step of base assembly by restricting the incorporation of the Hsm3-Rpt1-Rpt2-Rpn1 module, absence of such a checkpoint by Nas2 should increase base assembly. This aspect of Nas2 action could not be directly tested during endogenous base assembly, since the base continuously proceeds to the proteasome holoenzyme (lid-base-CP complex) by associating with the lid and CP. Since both lid and CP are absent in *E. coli*, the base does not progress into higher-order complexes and instead accumulates as a final complex, allowing us to test this prediction.

To abolish the proposed Nas2 steric hindrance, we silenced Nas2 expression in the heterologous system by introducing a premature stop codon into Nas2 (Fig. 2A, lane 2, labeled as *nas2D*). However, silencing Nas2 led to a defect in synthesis or stability of its cognate Rpt5, as seen from the truncated Rpt5 protein (Fig. 2A, lane 2). Such a phenomenon is not observed in yeast without Nas2 (30). Also, in a simpler version of the heterologous system co-expressing only Rpt4 and Rpt5, Rpt5 can stably exist irrespective of Nas2, forming the Rpt5-Rpt4 module (Fig. 2B). When all 9-base subunits are co-expressed in the heterologous system, Nas2 may become crucial for synthesis or stability of Rpt5. As an alternative approach to abolish Nas2 steric hindrance, we disrupted only the binding of Nas2 to Rpt5, by deleting a portion of Nas2 binding site on the Rpt5 C-terminal tail (the last 5 amino acids, indicated as *rpt5-Δ5*) (29,31). We confirmed that both Rpt5 and Nas2 are expressed normally in the *rpt5-Δ5* mutants, as in the wildtype cells (Fig. 2A, lanes 3, 4).

**Figure 2.**
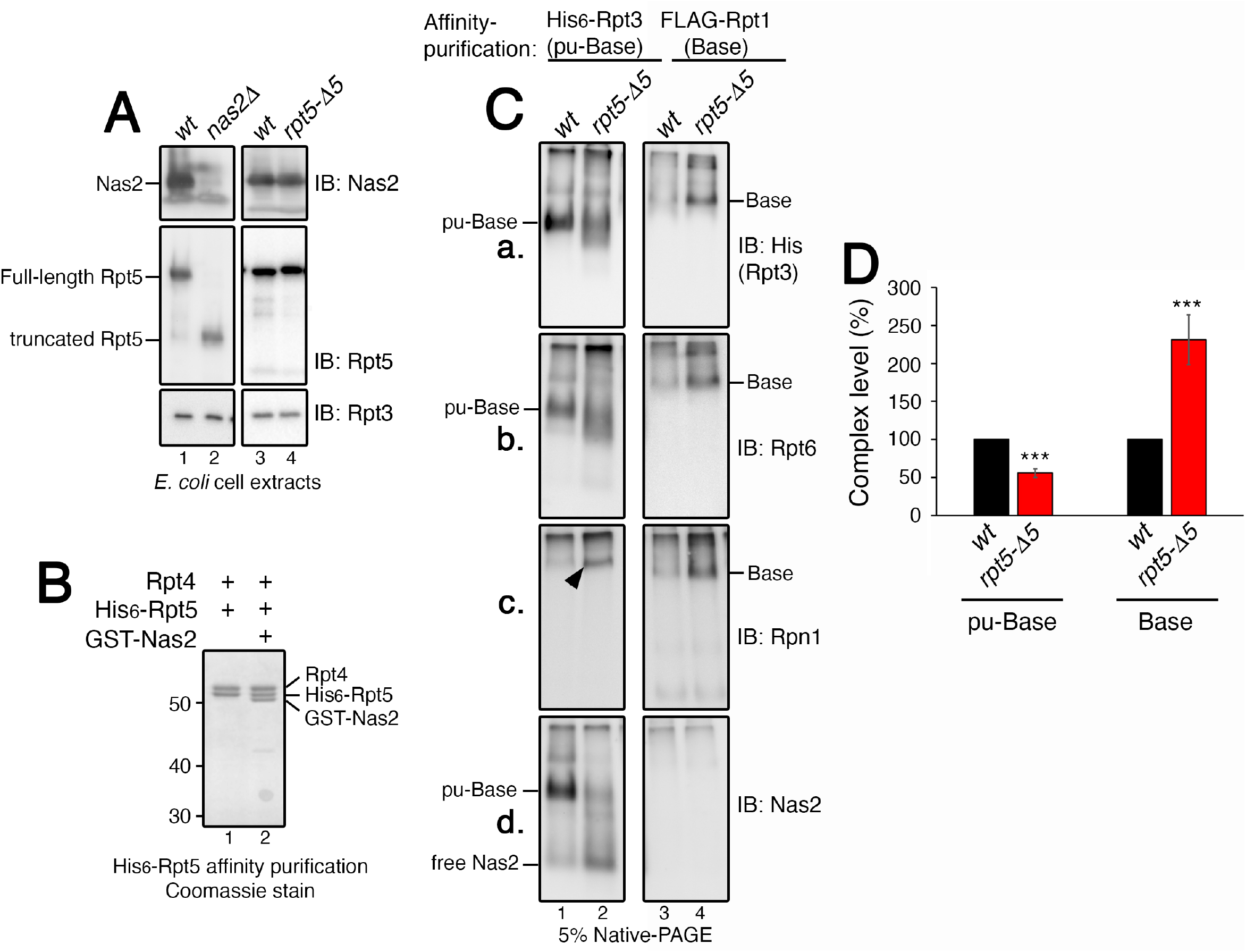
Nas2 blocks the premature progression of the penultimate base into the base. ***A***, Validation that both Nas2 and Rpt5 are expressed normally in the *rpt5-Δ5* mutant, which specifically disrupts Nas2 binding to Rpt5 (31). Nas2 silencing (*nas2Δ*) leads to an artifact, Rpt5 truncation, in the *E. coli* heterologous system co-expressing all 9 subunits of the base and 4 assembly chaperones. Whole cell extracts (15 μg) from the *E. coli* cells were analyzed by 10% Bis-Tris SDS-PAGE followed by immunoblotting for the indicated proteins. Rpt3, a subunit of the base and a loading control. ***B***, Rpt5 stably forms the Nas2-Rpt5-Rpt4 module in *E. coli* cells co-expressing two Rpt subunits, Rpt4 and His6-tagged Rpt5, with or without Nas2. Affinity purified protein (5 μg) using His6-tagged Rpt5 as a bait was subjected to 10% Bis-Tris SDS-PAGE and Coomassie stain. ***C***, The penultimate base more readily progresses into the base, upon disruption of Nas2 binding to the penultimate base, as in the *rpt5-Δ5* mutant. Affinity-purification was conducted in parallel, using His6-tagged Rpt3 and FLAG-tagged Rpt1, to obtain the penultimate base (pu-Base) and the base, respectively. Affinity-purified proteins (4 μg) were subjected to 5% native-PAGE and immunoblotting for indicated subunits of the base (Rpt3, Rpt6 and Rpn1) and Nas2. ***D***, Quantification of the relative levels of the penultimate base (pu-Base) and the base, using the data as in ***C***, panels [**a**] and [**b**] (mean ± SD, n=3 biological replicates, ***, p < 0.0005).

We then isolated the penultimate base via His6-Rpt3 affinity-purification from the heterologous system. We confirmed that Nas2 binding to the penultimate base was substantially reduced in the *rpt5-Δ5* mutant, as compared to its wild-type counterpart (Fig. 2C, [d], compare lane 2 to 1). Some free Nas2 is detected towards the bottom of the native gel, likely due to dissociation of residual Nas2 from the penultimate base (Fig. 2C, [d], lane 2). As compared to wild-type, the yield of the penultimate base in the *rpt5-Δ5* mutant was decreased to 50% of wildtype (Fig. 2C, [a, b], compare lane 1 to 2; see quantification in Fig. 2D). Some smeary appearance of the penultimate base complex might indicate more heterogenous conformations due to loss of Nas2 binding in this complex, since protein mobility on a native gel is influenced by not only its molecular mass but also conformation (32,33).

Notably, a decrease in the yield of the penultimate base in the *rpt5-Δ5* mutants is accompanied by a corresponding increase in the yield of the base complex, as seen from immunoblotting for representative base subunits, Rpt3, Rpt6, and Rpn1 (Fig. 2C, lane 4 in [a, b, c]; see quantification in Fig. 2D). A two-fold increase in the level of the base fits with the observed two-fold decrease in the level of the penultimate base in the *rpt5-Δ5* mutants (Fig. 2D). This result supports that the progression of the penultimate base into the base is accelerated without Nas2. Also, the penultimate step of base assembly in the *rpt5-Δ5* mutants already incorporated Rpn1, exhibiting Rpn1 signal in a slowly migrating complex (Fig. 2C, [c], arrowhead in lane 2). Given that Rpn1 is normally absent in the penultimate base (Fig. 1, lane 2), this result suggests that Rpn1 prematurely incorporated at the penultimate step of base assembly in the *rpt5-Δ5* mutants. Thus, without Nas2, the penultimate base can proceed to the base more readily. These data provide experimental evidence supporting the proposed model, in which Nas2 restricts the progression of the penultimate step to the final step for proper completion of the base.

### The penultimate base forms during endogenous base assembly in a Nas2-dependent manner

Based on our data supporting Nas2 action as a checkpoint for base assembly in the heterologous system, we sought to investigate whether and how Nas2 may provide a checkpoint during endogenous base assembly in yeast, *S. cerevisiae*. We first tested whether the endogenous penultimate base can be captured, since it has not been readily detectable as a discrete complex on a native gel. For this, we used the established strain harboring GFP-3xFLAG tagged Rpt6 (17,34). Since this strategy has been used to track Rpt6 incorporation into late-stage assembly intermediates, such as the base and RP (17,34), it should be able to also track the early segment of base assembly—the penultimate base and the preceding Rpt6-Rpt3 module (Fig. 3A). Following affinity-purification using the 3xFLAG tag on Rpt6, GFP fluorescence readily visualized Rpt6 incorporation into more abundant late-stage assembly intermediates, base and RP, and then into proteasome holoenzymes, which migrate closely together towards the top of the gel (Fig. 3B, [b]); this segment of base assembly into the proteasome holoenzyme is well established (17,34).

**Figure 3.**
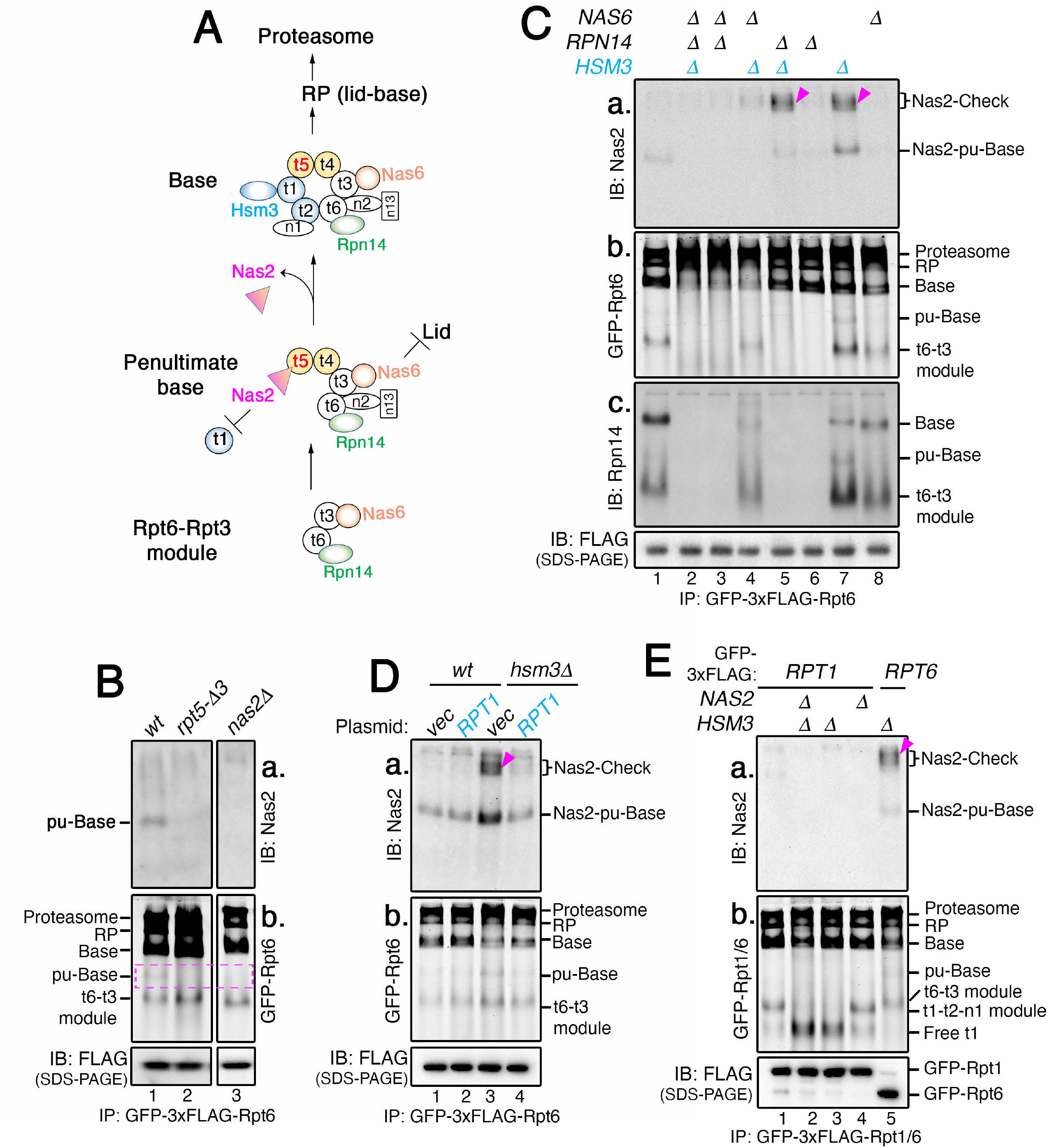
Nas2 activates a checkpoint during endogenous base assembly, when Rpt1 cannot be properly incorporated. ***A***, Cartoon showing the sequence of chaperone-mediated base assembly. Six distinct Rpt subunits (indicated as t1 through t6) undergo stepwise assembly into the base complex. The Rpt6-Rpt3 module with 2 chaperones, Rpn14 and Nas6, forms first, and then associates with the Rpt5-Rpt4 module harboring the Nas2 chaperone. Two non-ATPase subunits, Rpn2 and Rpn13 (n2 and n13) are then added (1,2). This penultimate base complex may utilize steric effects of Nas2 against Rpt1 (25) and of Nas6 against the lid (20,28). The base complex completes upon incorporation of the Rpt1-Rpt2-Rpn1 module via Hsm3, and release of Nas2 (25,26). For simplicity, this cartoon focuses on chaperone actions during early-stage base assembly, and does not show chaperone actions during the progression of the base into a proteasome holoenzyme. ***B***, The endogenous penultimate base forms in a Nas2-dependent manner. To capture the penultimate base and the preceding Rpt6-Rpt3 module in ***A***, affinity-purification was conducted using Rpt6 harboring a GFP-3xFLAG tag. Affinity-purified proteins (7.5 μg) were subjected to 5% native-PAGE, followed by immunoblotting for Nas2 [**a**] and Typhoon imaging for detection of GFP fluorescence on Rpt6 [**b**]. Anti-FLAG immunoblot was used a loading control for FLAG pulldown in ***B***, ***C***, ***D***, and ***E*** in Fig. 3. ***C***, Nas2 activates an assembly checkpoint, upon disruption of Hsm3 activity during endogenous base assembly. Nas2-Check complexes (arrowheads) are detected via affinity-purification using Rpt6 with GFP-3xFLAG tag. Affinity-purified proteins (7.5 μg) were subjected to 5% native-PAGE, followed by immunoblotting for Nas2 and Rpn14 [**a, c**] and by detection of GFP fluorescence to visualize Rpt6-containing complexes [**b**]. ***D***, Nas2-Checkpoint can be neutralized by compensating for disruption of Hsm3 activity, through Rpt1 overexpression. Empty vector (vec) or low-copy expression plasmid harboring *RPT1* was introduced into wild-type and *hsm3Δ* cells. Experiments were conducted as in ***B***. ***E***, Rpt1 cannot incorporate into the Nas2-Check complex, due to steric hindrance of Nas2 against Rpt1. Experiments were conducted as in ***B***, using the pulldowns with GFP-3xFLAG-tagged Rpt1 (lanes 1-4) and GFP-3xFLAG-tagged Rpt6 (lane 5). Lane 5 provides a positive control for the Nas2-Check complex (arrowhead) showing incorporation of Rpt6 into this complex.

Importantly, the endogenous penultimate base can be detected, albeit at a low level, migrating below the base in wild-type cells (Fig. 3B, see pu-Base, lane 1 in [a, b]); its subunit composition was validated by a proteomics analysis (Table S2). The existence of the penultimate base depends on its key component, the Nas2 chaperone, since it is no longer detectable upon disruption of Nas2 binding to the Rpt5 C-terminus, or deletion of *NAS2* (Fig. 3B, lanes 2, 3 in [a, b]; *rpt5-Δ3* lacks the last 3 amino acids as in ref. (35)). We also captured a step preceding the penultimate base, the Rpn14-Rpt6-Rpt3-Nas6 module, which is the fastest-migrating complex on a native gel (Fig. 3B, [b], t6-t3 module; see Table S2). In both *rpt5-Δ3* and *nas2Δ* cells, a decrease in the penultimate base was accompanied by the corresponding increase in the preceding Rpt6-Rpt3 module (Fig 3B, [b], lanes 2, 3), suggesting that Nas2 is needed for stable incorporation of this module into the penultimate base.

Both the Rpt6-Rpt3 module and the penultimate base complex are detected at a low level, supporting a view that these complexes progress rapidly through base assembly and into the proteasome holoenzyme (17,36). Also, both of these complexes are resolved better via a 5% native gel as used in our experiments, than the more commonly used 3.5% for separating larger proteasomal complexes. These two features explain why these complexes may have not been readily detectable in previous studies.

### The penultimate base exists as a part of the ordered assembly of the base via chaperones

We examined how this poorly characterized, penultimate step of base assembly, might be influenced by the actions of the other chaperones, which are proposed to act together with Nas2 for proper assembly of the base complex (Fig. 3A) (12,13,17,18,30). For this, we left Nas2 intact, since it is required for the penultimate base complex (Fig. 3B, lane 3), and deleted the other chaperones, *NAS6, RPN14*, and *HSM3*, individually or in combination. We then followed the Rpt6-Rpt3 module and its progression into the penultimate base, by conducting GFP-3xFLAG-Rpt6 affinity-purification and native gel analysis.

Whenever the Rpn14 chaperone in the Rpt6-Rpt3 module (Fig. 3A) was deleted alone, or together with another chaperone, this module was no longer detectable with little to no penultimate base (Fig. 3C, [b, c], lanes 2, 3, 5, 6). This result suggests that Rpn14 facilitates formation of the Rpt6-Rpt3 module and its progression into the penultimate base, explaining Rpn14 action during early-stage base assembly (34). By comparison, Nas6, the other chaperone in the Rpt6-Rpt3 module (Fig. 3A), seems to be more important for the penultimate base than the module itself. In the *nas6Δ* single mutants, the penultimate base was not readily detectable although the Rpt6-Rpt3 module still exists (Fig. 3C, lane 8 in [b, c]). This result can be attributed to the known activity of Nas6, which sterically prevents a premature base complex, such as the penultimate base, from associating with the lid or CP (Fig. 3A) (16,20–22,28). Without Nas6, the penultimate base may prematurely proceed into the proteasome holoenzyme (lid-base-CP complex), depleting the cellular pool of penultimate base. These results suggest that the penultimate step of base assembly relies on proper actions of not only Nas2 (Fig. 3B), but also Rpn14 and Nas6, which incorporate into this step from the preceding Rpt6-Rpt3 module.

To complete the base complex, the penultimate base must associate with the incoming Rpt1-Rpt2-Rpn1 module via the Hsm3 chaperone (Fig. 3A). Without Hsm3, the Rpt1-Rpt2-Rpn1 module cannot form efficiently (see Fig. 3E, lanes 2, 3 in [b]) (12,14,16,17). This phenomenon explains some accumulation of both the penultimate base and Rpt6-Rpt3 module in *hsm3Δ* cells (Fig. 3C, [b, c], lane 7), since these complexes cannot properly progress into the base, without the incoming Rpt1-Rpt2-Rpn1 module. When *HSM3* is deleted together with *RPN14* or *NAS6*, which are needed for the penultimate base itself, little to no penultimate base was detectable (Fig. 3C, [b, c], lanes 2, 4, 5).

Taken together, our data demonstrate that the penultimate step of base assembly via Nas2 (Fig. 3B) is integrated with the actions of the other chaperones (Rpn14, Nas6 and Hsm3) (Fig. 3C). The steric effect of Nas2 may restrict the progression of the penultimate base, to ensure its proper subunit composition together with Rpn14 and Nas6, until the incorporation of the last module by Hsm3.

### Nas2 activates a checkpoint during endogenous base assembly, when Rpt1 cannot be properly incorporated

Combined with our data that the penultimate step of base assembly relies on Nas2 (Figs. 2C, 3B), the current model proposes that completion of the base complex is accompanied with release of Nas2, due to its steric conflict with the incoming Rpt1 at the final step of base assembly (25–27). This model predicts that, if this final step of base assembly is defective, Nas2 may not release, as a mechanism to provide a checkpoint to prevent formation of a defective base complex.

Indeed, Nas2 can be found, not only in the penultimate base, but also in additional slow-migrating complexes, prominently in *hsm3Δrpn14Δ* cells and *hsm3Δ* cells (Fig. 3C, [a], see arrowheads in lanes 5, 7). This result suggests that these additional Nas2-containing complexes form, due to disruption of Hsm3 activity. Hsm3 is responsible for adding the final, Rpt1-Rpt2-Rpn1 module to the penultimate base, triggering Nas2 release for completion of the base (Fig. 3A). Without Hsm3, these additional Nas2-containing complexes may form due to disruption of Nas2 release, since the penultimate base cannot properly progress into the final step of base assembly. We will refer to this complex as Nas2-Check henceforth. Nas2-Check complexes may form at a low-level, migrating at a similar position as RP as a more diffuse signal (Fig. 3C, lanes 5, 7 in [a]).

Nas2-Check complexes form upon deletion of *HSM3*, but become nearly undetectable when *NAS6* was also deleted, as in *nas6Δrpn14Δhsm3Δ* and *nas6Δhsm3Δ* cells (Fig. 3C, [a], lanes 2, 4). This result suggests that the Nas2-Check complex relies on the steric effect of Nas6, as seen in the penultimate base complex (Fig. 3C, [b, c], lane 8). Nas6 may shield such a premature base complex by obstructing its association with the incoming lid or CP until base assembly is complete (16,20–22,28). In support of this idea, when *NAS6* is deleted together with at least one or more chaperones, the cellular pool of base is noticeably depleted (Fig. 3C, [b], lanes 2, 3, 4; see base). We also confirmed that Nas2 itself is required for the existence of the Nas2-Check complex, since this complex is no longer detected upon deletion of *NAS2*, as seen in *nas2Δ hsm3Δ* cells (Fig. S2, compare lane 3 to 2). Thus, the Nas2-Check complex requires steric effects of both Nas2 and Nas6, arising specifically upon disruption of Hsm3 activity during base assembly.

If the Nas2-Check complex forms as a potential checkpoint upon loss of Hsm3 activity, this checkpoint should be neutralized upon compensating for the loss of Hsm3 activity. Since Hsm3 helps recruit Rpt1 into the final step of base assembly (Fig. 3A), we tested whether increasing Rpt1 expression might be sufficient to promote this final step, even without Hsm3 (16). For this, we introduced a low-copy Rpt1 expression plasmid into wild-type and *hsm3Δ* cells (Fig. 3D). In wild-type cells, there was no discernable effect on base assembly upon increased Rpt1 expression (Fig. 3D, [a, b], lanes 1, 2). Notably, in *hsm3Δ* cells, the Nas2-Check complex was abolished upon increased Rpt1 expression (Fig. 3D, [a], compare lane 3 to 4). Also, the level of the base complex in *hsm3Δ* cells became more comparable to that in wild-type (Fig. 3D, [b], compare lane 4 to 1). Thus, if Rpt1 can efficiently incorporate at the final step of base assembly, Nas2 may release normally, allowing for completion of base assembly. These results explain the basis for the Nas2-dependent checkpoint, that Nas2 may distinguish normal vs. defective base assembly, depending on whether Rpt1 can incorporate properly into the penultimate base complex.

Next, we tested whether the Nas2-dependent checkpoint utilizes the steric hindrance of Nas2 against Rpt1 incorporation, thereby blocking completion of the base complex (25). For this, we tested whether the Nas2-Check complex can be detected in the same complex with Rpt1 during base assembly. We conducted affinity-purification using Rpt1 harboring GFP-3xFLAG tag upon deletion of *HSM3* and *NAS2* individually and together. In wild-type cells, Rpt1 incorporated into the Rpt1-Rpt2-Rpn1 module, and then into the base, RP, and finally into the proteasome holoenzyme, as detected by Rpt1 GFP fluorescence (Fig. 3E, [b], lane 1). Upon deletion of *HSM3*, free Rpt1 becomes prominent, since it cannot efficiently form the Rpt1-Rpt2-Rpn1 module without Hsm3 (Fig. 3E, [b], lanes 2, 3) (12,14,16,17). Importantly, the Nas2-Check complex was not detectable in any of Rpt1-containing complexes isolated from *hsm3Δ* cells, although it is readily detectable as a part of Rpt6-containing complexes in *hsm3Δ* cells (Fig. 3E, [a], compare lane 3 to 5). This result demonstrates that the Nas2-Check complex cannot incorporate Rpt1, supporting our conclusion that the Nas2-dependent checkpoint utilizes the steric effect of Nas2 against Rpt1, to block formation of a defective base complex.

### The Nas2-dependent checkpoint arises from the penultimate step of base assembly

As a readout for checkpoint activity during base assembly, the Nas2-Check complex migrates more slowly on native gels than the penultimate base, which is normally the last step harboring Nas2 during base assembly (Fig. 3C-E). We sought to examine whether the Nas2-Check complex arises from the penultimate base. For this, we specifically tracked Nas2-containing complexes, using Nas2 itself as a bait for affinity-purification in our panel of chaperone deletion strains.

Except the penultimate base, Nas2-Check complexes are the only additional higher-order complex as evident from the native gel, and they are detected prominently in *rpn14Δhsm3Δ* and *hsm3Δ* cells (Fig. 4A, lanes 5, 8; arrowheads). These results are consistent with those in our experiments using Rpt6 as a bait for tracking the Nas2-Check complex during base assembly (Fig. 3C, lane 5, 7; arrowheads). These data together support that the Nas2-Check complex arises directly from the penultimate step of base assembly. We noticed that the Nas2-Check complexes were detected as doublets (Fig. 4A, lanes 5, 8, see an additional low-intensity signal below the arrowhead), suggesting that some small fraction of the Nas2-Check complexes might potentially exist in an additional form. Since the Nas2-Check complexes depend on the steric effect of Nas6 (Fig. 3C, [a], lanes 2, 4), they become nearly undetectable in *nas6Δrpn14Δhsm3Δ* and *nas6Δhsm3Δ* cells (Fig. 4A, lanes 2, 4); in these samples, Nas2 exists mostly in a Nas2-Rpt5-Rpt4 module and free Nas2. On the other hand, when Nas2-Check complex forms as in *rpn14Δhsm3Δ* and *hsm3Δ* cells, the Nas2-Rpt5-Rpt4 module and free Nas2 are decreased comparatively (Fig. 4A, lane 5, 8). Since the Nas2 chaperone is normally recycled by releasing from the completed base complex, disruption of its release due to Nas2-Check complex formation would interfere with such recycling to a new round of Nas2-dependent steps during base assembly.

**Figure 4.**
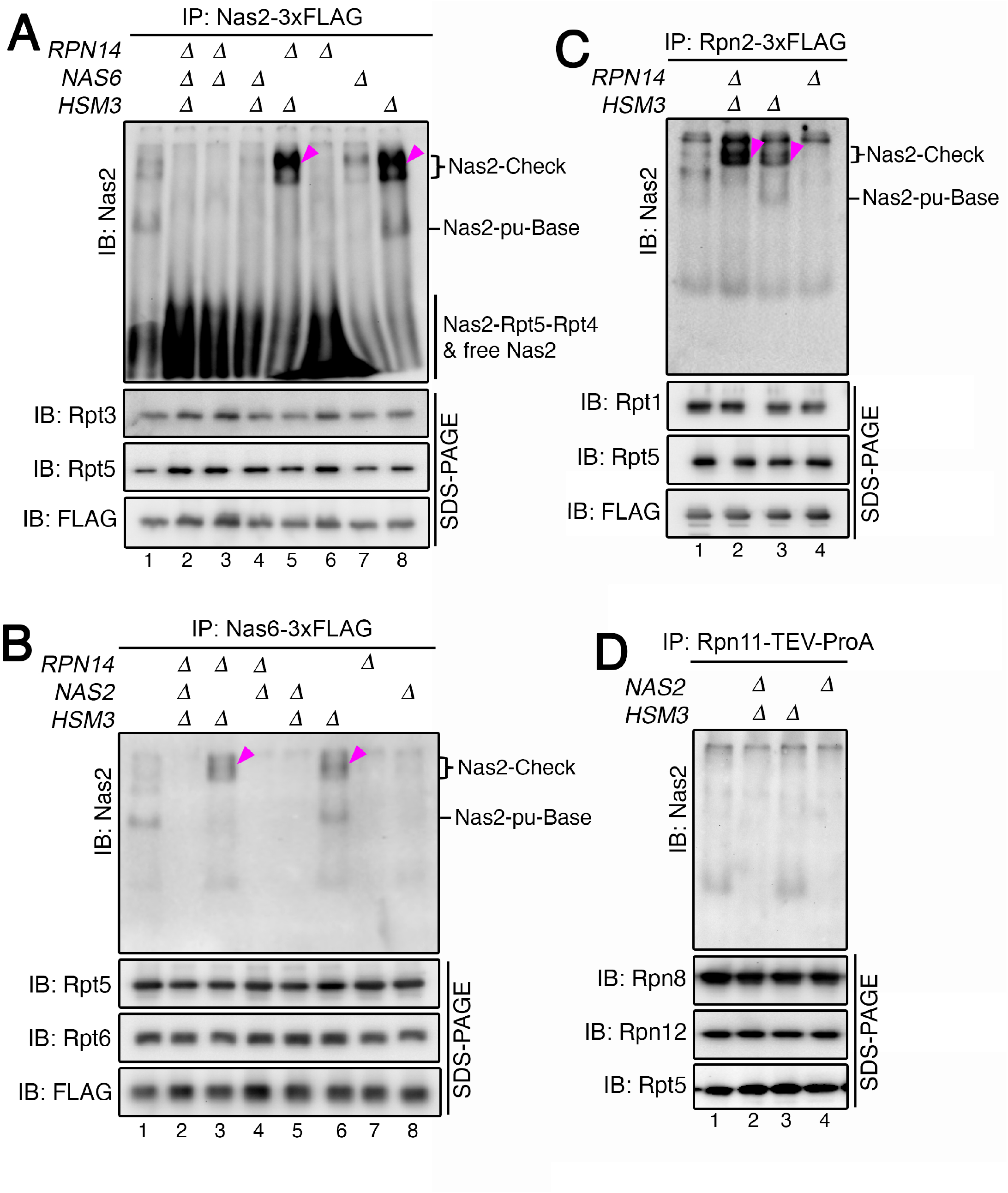
The Nas2-checkpoint arises from the penultimate step of base assembly. ***A***, Nas2-Check complexes (arrowheads) arise from the penultimate step of base assembly. Affinity-purification was conducted, using 3xFLAG tagged Nas2 as a bait. To detect the Nas2-Check complexes, Nas2 pulldown proteins (5 μg) were subjected to 5% Native-PAGE and immunoblotting for Nas2. To confirm that largely comparable amounts of the complexes were analyzed, pulldown proteins (1 μg) were also subjected to 10% Bis-Tris SDS-PAGE followed by immunoblotting for 3 controls: 2 representative Rpt subunits and each specific 3xFLAG-tagged bait protein for pulldowns (panels ***A***, ***B*** and ***C*** in Fig. 4). ***B, C***, Validation that the Nas2-Check complexes (arrowheads) arise from the penultimate base, using two distinct components: the Nas6 chaperone and a non-ATPase subunit, Rpn2. Nas2-Check complexes are readily detectable in the pulldowns with Nas6-3xFLAG (***B***) as well as Rpn2-3xFLAG (***C***), upon analyzing 2.5 and 7.5 μg of the pulldown proteins, respectively. Experiments were conducted as described in ***A***. ***D***, Nas2-Check complexes do not incorporate the lid. Lid-containing complexes were isolated via affinity-purification using a representative lid subunit, Rpn11, harboring TeV-ProA affinity tag. Pulldown proteins (7.5 μg) were subjected to 5% Native-PAGE and immunoblotting for Nas2 as in ***A***. To ensure that comparable amounts of the lid-containing complexes were analyzed, pulldown proteins (1 μg) were subjected to 10% Bis-Tris SDS-PAGE followed by immunoblotting for 2 additional lid subunits (Rpn8 and Rpn12) and a lid-base subunit (Rpt5).

Since our data suggest that the Nas2-dependent checkpoint utilizes the steric activity of Nas6 (Figs. 3C and 4A, lanes 2, 4), we tested whether the Nas2-Check complex contains Nas6, arising together from the penultimate base, in which both Nas2 and Nas6 exist (Fig. 3A). Indeed, the Nas2-Check complex was readily detected in the Nas6 pulldown, specifically in *rpn14Δhsm3Δ* and *hsm3Δ* cells (Fig. 4B, lanes 3, 6, see arrowheads). This result supports that the Nas2-Check complex contains Nas6, and the steric effect of Nas6 helps maintain this Nas2-dependent checkpoint. When *NAS2* was additionally deleted in these cells, the Nas2-Check complex was no longer detectable (Fig. 4B, lanes 2, 5), confirming that the Nas2-Check complex also depends on the steric effect of Nas2 itself.

Based on the proposed order of base assembly, Rpn2, a non-ATPase subunit, is recruited into the penultimate base later than the ATPase subunits, such as Rpt6 (Fig. 3A) (26,27). We tested whether Rpn2 pulldowns also include the Nas2-Check complex. When we conducted affinity-purification using Rpn2 as a bait, the Nas2-Check complex is detected in both *rpn14Δhsm3Δ* and *hsm3Δ* cells (Fig. 4C, lane 2, 3, see arrowheads). This result suggests that the Nas2-Check complex is likely to contain all subunits in the penultimate base complex. As expected from the activity of Nas6, that it obstructs binding of the lid (20,28), the Nas2-Check complex was not found in the affinity-purification using Rpn11, a representative subunit of the lid (Fig. 4D).

Taken together, these data suggest that the Nas2-dependent checkpoint arises from the penultimate step of base assembly. The steric activity of Nas2 may arrest the progression of the penultimate base, upon any disruption of Hsm3 activity for the incoming Rpt1-Rpt2-Rpn1 module, to prevent formation of a defective base complex.

### Nas2 recognizes proper completion of base assembly via ATP hydrolysis by Rpt4

The current model explains how Nas2 sterically hinders Rpt1 addition at the checkpoint, as shown by our data (Figs. 1, 2C, and 3C-E) (25), but does not explain how this same steric effect may also trigger Nas2 release, allowing Rpt1 addition to complete the base complex. This aspect is important for understanding how this checkpoint can be relieved via Nas2 release, suggesting that an additional feature might regulate the steric effect between Nas2 and Rpt1 during base assembly. We hypothesized that such a feature might be the fundamental activity of the assembled base— ATP hydrolysis, since it is initiated upon completion of the heterohexameric Rpt ring in the base (18) and is known to cause conformational changes in its subunits (6–9). To test this hypothesis, we fixed individual Rpt proteins in an ATP-bound state by using *rpt-EQ* mutants; glutamine substitution for the conserved Walker B glutamate blocks ATP hydrolysis by a given Rpt protein (22,37). Since the *rpt1-EQ, rpt4-EQ* and *rpt5-EQ* mutants are lethal in yeast (37), they were expressed using plasmids in the presence of their endogenous chromosomal *RPT* gene to maintain cell viability. We predicted that the Nas2-check complex should form if ATP hydrolysis by any specific ATPase is necessary for Nas2 release and Rpt1 incorporation.

Specifically, inhibition of ATP hydrolysis by Rpt4 resulted in the formation of Nas2-Check complexes (Fig. 5A, [a], lane 3; see arrowhead). Although Nas2 directly binds to Rpt5, a direct neighbor of Rpt4 during heterohexameric Rpt ring assembly (Fig. 3A), Nas2 released normally in the *rpt5-EQ* mutants (Fig. 5A, [a], lane 7). This result suggests that Nas2 release is not influenced by the nucleotide state of its cognate Rpt5, but by its direct neighbor Rpt4. In the *rpt5-EQ* background, the Nas2-Check complex formed only when *HSM3* was deleted as in *hsm3Δ rpt5-EQ* double mutants (Fig. 5A, [a], lane 8). Nas2 also released normally in *rpt1-EQ, rpt2-EQ, rpt3-EQ* and *rpt6-EQ* mutants, as evidenced by no detectable Nas2-Check complex formation (Fig. 5B, lanes 2, 4, 6, 8). These results suggest the specificity of ATP hydrolysis by Rpt4, as a requirement for Nas2 release for proper completion of the base complex.

**Figure 5.**
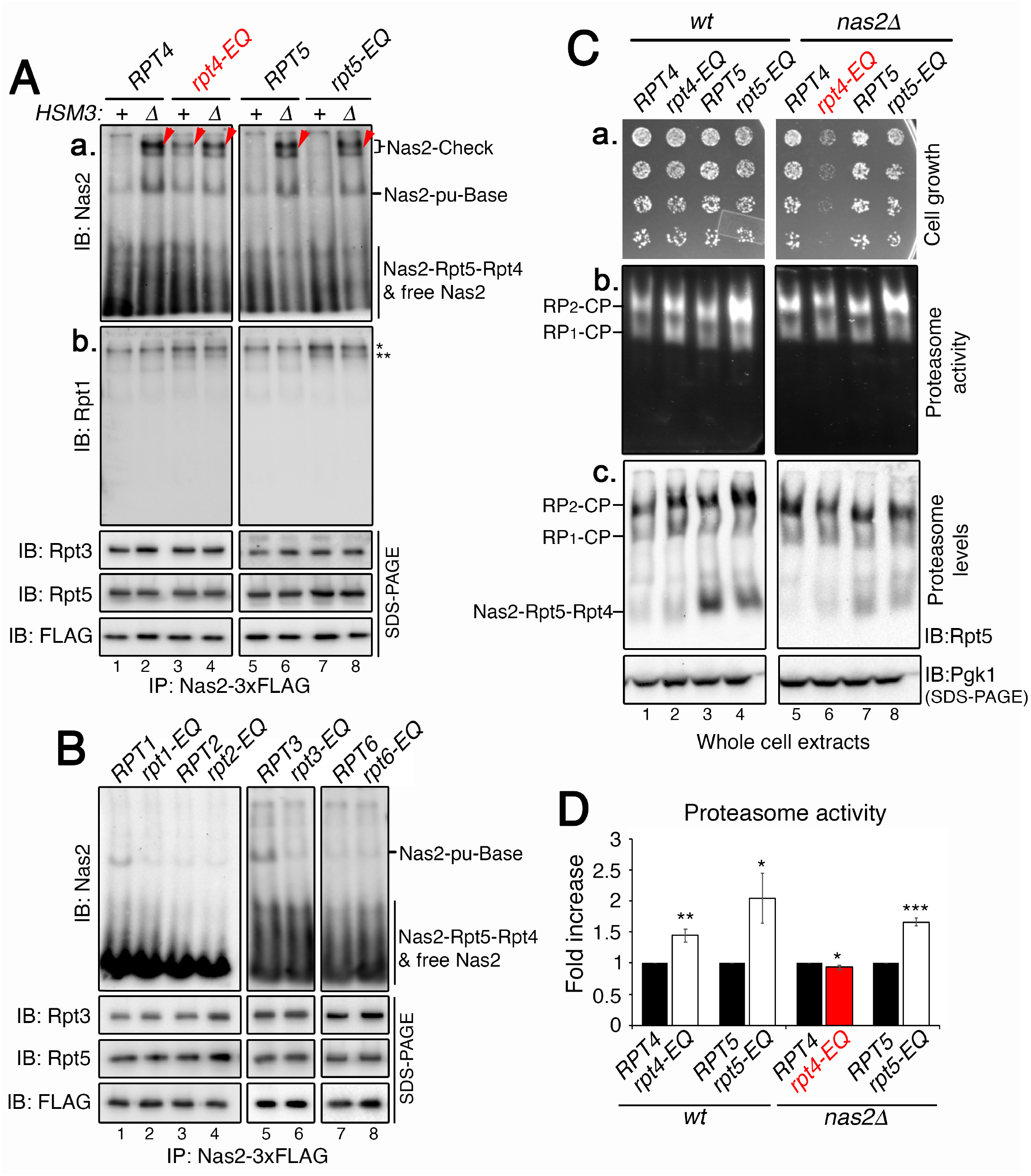
Nas2 recognizes proper completion of base assembly via ATP hydrolysis by Rpt4. ***A, B***, Nas2-dependent checkpoint is activated, specifically upon inhibiting ATP hydrolysis by Rpt4. Affinity-purification was conducted using 3xFLAG-tagged Nas2. The purified proteins (5 μg) were subjected to 5% native-PAGE followed by immunoblotting for indicated proteins (*, top of the resolving gel; **, background signals that exist irrespective of the Nas2-Check complex formation, for example, in lanes 7 and 8 in [b]). To ensure that largely comparable amounts of the complexes were analyzed, pulldown proteins (4 μg) were also subjected to 10% Bis-Tris SDS-PAGE and immunoblotting for 2 representative Rpt subunits and the 3xFLAG-tagged Nas2 itself, as a bait for pulldowns. ***C***, The Nas2-dependent checkpoint ensures efficient assembly of proteasome holoenzymes and cell growth upon heat stress. To assess heat sensitivity of the indicated yeast strains, 3-fold serial dilutions of yeast cells were spotted onto synthetic dropout media lacking leucine (a selection marker for the indicated plasmids) and were incubated at 37°C for 2-3 days [**a**]. Proteasome activity and level were analyzed using yeast cultures upon heat stress at 37°C for 3.5 hrs (see Experimental Procedures for details). Whole cell extracts (50 μg) were subjected to 3.5% native-PAGE and in-gel peptidase assays [**b**], followed by immunoblotting [**c**]. The Nas2-Rpt5-Rpt4 module is more noticeable in lanes 3, 4 [**c**] since the total abundance of Rpt5 is increased due to combined plasmid-born and chromosomal expression of *RPT5* allele, enhancing Rpt5 binding to its cognate Nas2 chaperone for the formation of this module. Pgk1, loading control. ***D***, Quantification of proteasome activity (RP2-CP) from data as in ***C***, panel [**b**]. Fold increase of the proteasome activity in *rpt4-EQ* and *rpt5-EQ* was calculated, relative to that of their wild-type counterpart (mean ± SD, n=3 biological replicates, *, p < 0.05, **, p < 0.005, ***, p < 0.0005).

The requirement of Rpt4 ATP hydrolysis for Nas2 release reveals how Nas2 may recognize proper completion of base assembly. Nas2 specifically binds to Rpt5, which is positioned directly between Rpt4 and Rpt1, arranged as in Rpt4-Rpt5-Rpt1 in the heterohexameric Rpt ring of the base complex (Fig. 3A). As seen in *rpt4-EQ* cells, inhibiting Rpt4 ATP hydrolysis alone can disrupt Nas2 release, although Hsm3 remains intact to recruit Rpt1 (Fig. 5A, [a], lane 3). This result suggests that Hsm3 activity alone may not be sufficient for ensuring both proper Rpt1 incorporation and Nas2 release, but additionally requires Rpt4 ATP hydrolysis for these events. If Rpt4 ATP hydrolysis is required only for Nas2 release, but not for Rpt1 addition, Nas2 should be found in the base complex with Rpt1 in it. However, Rpt1 does not appear to incorporate into any of the Nas2-containing complexes since Rpt1 is detected only at a background level in these complexes (Fig. 5A, [b], lane 3), suggesting proper Rpt1 incorporation into the final step of base assembly may also depend on Rpt4 ATP hydrolysis.

Taken together, during heterohexameric Rpt ring assembly of the base, Nas2 may recognize Rpt4 ATP hydrolysis on one side of Rpt5, and Rpt1 incorporation to the other side of Rpt5. Upon proper completion of these two events on both sides of Rpt5, Nas2 may release, thereby ensuring correct completion of the base.

### The Nas2-dependent checkpoint ensures proper assembly of the proteasome holoenzyme

We examined to what extent the Nas2-dependent checkpoint may influence proteasome holoenzyme assembly in stressed conditions, such as heat, in which misfolded proteins are generated in a large quantity and require proteasome-mediated degradation for cell survival. When Nas2 is intact, both *rpt4-EQ* and *rpt5-EQ* cells tolerated heat stress at 37°C, similarly to their corresponding control (Fig. 5C, [a], compare lane 2 to 1, and 4 to 3). Proteasome activities and levels were increased in both *rpt4-EQ* and *rpt5-EQ* cells by 1.5- to 2-fold, due to a compensatory mechanism that raises proteasome gene expression upon deficit in proteasome function, such as defects in ATP hydrolysis as in these mutants (Fig. 5C, [b, c], compare lane 2 to 1, and 4 to 3; see Fig. 5D for quantification) (38–40).

Upon disabling the Nas2-dependent checkpoint by deletion of *NAS2*, the *rpt4EQ* cells could not tolerate the heat stress, unlike the *rpt5-EQ* cells (Fig. 5C, [a], lane 6 vs. 8). This growth defect of *rpt4-EQ nas2Δ* double mutant cells is due to a failure to increase proteasome holoenzyme assembly, since both the activity and level of the proteasome holoenzymes remained similar between the *rpt4EQ nas2Δ* double mutants and their corresponding control (Fig. 5C, [b, c], compare lane 6 to 5; see quantification in Fig. 5D). On the other hand, in the *rpt5-EQ nas2Δ* double mutants, the proteasome holoenzyme level was increased by 1.7-fold, similarly to the extent seen between the *rpt5-EQ* single mutant and its corresponding control (Fig. 5C, [b, c], compare lane 8 to 7, and lane 4 to 3; see quantification in Fig. 5D). These results support that the Nas2-dependent checkpoint is specifically connected with Rpt4 ATP hydrolysis during base assembly, providing a mechanism crucial for proper and efficient formation of the proteasome holoenzyme.

## DISCUSSION

### Nas2 monitors dual requirements for completion of base assembly

In the present study, we describe chaperone-mediated regulation at one of the least characterized steps of proteasome assembly, the completion of the base complex from the penultimate base. This step needs to ensure whether the 9-subunit base formed properly, prior to its further association with the other sub-assemblies, lid and CP, for proteasome holoenzyme formation. To investigate this aspect of chaperone-mediated base assembly, we examined the Nas2 chaperone, based on the known correlation between Nas2 release and completion of the base complex (25–27).

Our findings reveal dual criteria by which Nas2 recognizes proper completion of the base complex: 1) addition of the last module, Rpt1-Rpt2-Rpn1 via the Hsm3 chaperone, and 2) ATP hydrolysis by Rpt4 (Figs. 3–5). Through these two tightly connected criteria, Nas2 may ensure not only correct subunit composition, but also the fundamental activity of a newly assembled base. Such ability of Nas2 can be explained by its positioning, in that Nas2 directly binds to its cognate Rpt5, which exists between Rpt1 and Rpt4, as in Rpt1-Rpt5-Rpt4 in a heterohexameric Rpt ring (Fig. 3A). Thus, the Nas2 chaperone on Rpt5 can recognize whether Rpt1 is added properly to one side of Rpt5, based on ATP hydrolysis by Rpt4 on the other side of Rpt5 (Figs. 3, 5). Since ATP hydrolysis is initiated upon completion of the heterohexameric Rpt ring (18), only the proper incorporation of Rpt1 may activate ATP hydrolysis by Rpt4, perhaps as the first event of ATP hydrolysis in a newly assembled base. Conformational changes by Rpt4 ATP hydrolysis may then transmit to the directly neighboring Rpt5, as a signal to release Nas2. In this way, Nas2 can release, only upon correct completion of the base both compositionally and functionally.

Since it is well established that ATP hydrolysis by the base causes specific conformational changes (6–9), Rpt4 ATP hydrolysis may ensure proper base assembly not only by triggering Nas2 release, but also by contributing to conformational changes essential for protein degradation. This idea is supported by the evidence that the *rpt4-EQ* base exhibits non-productive conformational changes, abolishing the ability to degrade protein substrates by the proteasome holoenzyme, as seen from the recombinant base from the *E. coli* heterologous system (29). The *rpt4-EQ* mutation is lethal in yeast, further supporting the requirement of Rpt4 ATP hydrolysis for proteasome function (37). For example, the Rpt1-Rpt2-Rpn1 module, whose incorporation into the base is connected with ATP hydrolysis by Rpt4 (Fig. 5A), consists of subunits crucial for ubiquitin recognition and processing during protein degradation by the proteasome holoenzyme. Rpn1 is the major ubiquitin receptor, which recognizes polyubiquitinated protein substrates for the proteasome holoenzyme (41). Also, Rpn1 and Rpt1 together regulate the rate of protein degradation by controlling the activity of its bound Ubp6, a deubiquitinase of the proteasome (42,43). These activities of Rpt1 and Rpn1 are tightly coordinated via conformational changes by the base complex, supporting the importance of Nas2 action in ensuring that the newly assembled base can undergo proper conformational changes.

### The Nas2-dependent checkpoint contributes to sequential checkpoints for base assembly

Our findings suggest that Nas2 activates a checkpoint for base assembly, when one of its dual criteria for completion of base assembly is not satisfied, as seen in *hsm3Δ* and *rpt4-EQ* cells (Figs. 3–5). At this checkpoint, Nas2 remains on the penultimate base, instead of releasing from it and allowing its progression into the base. The Nas2-dependent checkpoint complex can be distinguished from the normal, Nas2-bound penultimate base, due to its slower mobility on native gels (Fig. 3–5), which might result from different composition or conformation, or both. Although the precise composition of the Nas2-Check complex remains to be determined, it appears to lack the same Rpt1-Rpt2-Rpn1 module (Figs. 3E, 5A), like the penultimate base, supporting that Nas2 continues to hinder the incorporation of these subunits at the checkpoint. Only 4 Rpt subunits exist in the penultimate base and they cannot yet form a stable ring-shaped complex. If such a complex cannot readily proceed into the base complex due to a defect, as is the case for the Nas2-Check complex, it might adopt a more open or flexible conformation, resulting in a slower mobility in the native gel.

In many cancer cells, a human ortholog of Hsm3, S5b, is found to be silenced (44), exhibiting a situation as in *hsm3Δ* cells in our experiments. S5b silencing is suggested to be responsible for altered proteasome activity in these cancer cells (44). A future study will be needed to determine whether the Nas2-dependent checkpoint can be properly activated in cancer cells, and how it might influence proteasome assembly and functions, since this Nas2-dependent checkpoint is needed to satisfy an increased demand for proteasome holoenzyme assembly (Fig. 5C, D).

An initial model of base assembly highlights that chaperones’ binding to ATPases sterically hinders the association of the base with the lid and CP, until base assembly is complete (15,16,19,23,25). This model raises an important question: how might chaperones recognize proper completion and progression of the base into the proteasome? Combined with previous findings (20,22,28), our present study can help address this question. Nas2 can specifically recognize the nucleotide state of Rpt4 as a requirement for proper completion of the base (Fig. 5A, B). As the base further matures into the proteasome holoenzyme, the other 3 chaperones (Rpn14, Hsm3 and Nas6) can individually distinguish the nucleotide state of specific Rpt subunits (20,22,28). These findings together propose a common feature of these chaperones, that each chaperone may recognize specific criteria for proper completion of a given assembly step, based on the nucleotide state of specific Rpt subunits. Steric effects of each chaperone may be regulated through this mechanism, to distinguish normal vs. defective assembly events. In this way, chaperones may provide a series of checkpoints, throughout the multi-step assembly process of the base and its incorporation into the proteasome holoenzyme.

## EXPERIMENTAL PROCEDURES

### Yeast Strains, Plasmids and Biochemical Reagents

A complete list of yeast strains and plasmids are provided in Tables S3 and S4, respectively. Unless specified otherwise, all yeast strains were cultured in YPD medium at 30°C. Yeast strains harboring specific gene expression plasmids were grown in synthetic dropout medium lacking the appropriate auxotrophic marker for the plasmids at 30°C. All yeast manipulations were conducted using standard procedures (45). Yeast strains harboring multiple engineered alleles were generated through crossing two appropriate strains, followed by sporulation and dissection. When the same markers were used for two alleles, PCR was used to distinguish strains with each of those two alleles.

At least two biological replicates were performed for all biochemical and imaging experiments. Details of biochemical reagents are included in each specific Experimental Procedure section. A complete list of antibodies is provided in the Table S5. All antibodies were used at 1:3000 dilutions except anti-Pgk1, which was used at 1:10,000.

### Affinity purification of the base and penultimate base from the heterologous *E. coli* system

*E. coli* BL21-star (DE3) cells were co-transformed with 3 plasmids, which were obtained from Andreas Martin’s laboratory (29): pCOLA-1 (FLAG-Rpt1, Rpt2, His6-Rpt3, Rpt4, Rpt5, Rpt6), pETDuet-1 (Rpn1, Rpn2, Rpn13), and pACYCDuet-1 (Nas2, Nas6, Hsm3, Rpn14). Genes for rare tRNAs were also included in the pACYCDuet-1 plasmid (29). In Fig. 2A (lane 2), Nas2 in the pACYCDuet-1 plasmid has a premature stop codon at amino acid 65. In Fig. 2A (lane 4) and Fig. 2C (lanes 2, 4), the pCOLA-1 plasmid contains Rpt5 with last 5 amino acids deletion. All plasmids are listed in Table S4. *E. coli* cells were grown in 35 mL LB media at 37°C overnight, and were inoculated into 2 L fresh LB media next day at OD_600_=0.05, and were grown to OD_600_ = 0.6. Cultures were cooled to room temperature, and were induced using 1 mM isopropyl-β-D-thiogalactopyranoside (IPTG) at 18°C in water bath overnight. Cells were harvested by centrifugation at 4000 x *g* for 10 min at 4°C. Cell pellets were washed once with 1 pellet volume of cold lysis buffer (50 mM Na-phosphate [pH 7.0], 300 mM NaCl, 10% glycerol], which is supplemented with 1 mM β-mercaptoethanol, and were centrifuged at 4000 x *g* for 10 min. Cell pellets were resuspended using the residual buffer. The resulting cell resuspension was frozen in a drop-wise manner into liquid nitrogen, and was ground using a mortar and pestle in the presence of liquid nitrogen. The ground cryo-powders were hydrated with 3 volumes of lysis buffer containing 1 mM β-mercaptoethanol, protease inhibitors and ATP (1 mM), on ice for approximately 20 min with intermittent vortexing. ATP (1 mM) was included for the rest of the purification procedure. Typically, 2 L cultures generate 15 mL cryo-powders, which are then hydrated with 45 mL of lysis buffer. Triton X-100 was added to 0.2% final to aid the solubilization of the proteins on ice for 10 min. Cleared lysates were obtained by centrifugation at 20,000 x *g* at 4°C for 30 min.

To isolate the penultimate base using His6-Rpt3 as a bait, the cleared lysates were incubated with 250 μL of TALON metal affinity resin (Clontech, 635502) for 1hr at 4°C. Beads-bound proteins were collected by centrifugation at 3000 rpm for 5 min, followed by 3 washes with 5 beads volume of lysis buffer containing 0.2% Triton X-100, using filter column. The penultimate base was eluted from the beads using 3 beads volume of lysis buffer containing 150 mM imidazole by rotating for 1 hr at 4°C.

To isolate the base complex using FLAG-Rpt1 as a bait, the cleared lysates were incubated with 250 μL of anti-FLAG M2-agarose beads (Sigma, A2220-5ML) for 2 hrs at 4°C. The beads were washed 3 times with 5 beads volume of proteasome buffer containing 150 mM NaCl. The base complex was eluted using 3 beads volume of proteasome buffer containing 0.2 mg/mL FLAG peptides (GLP Bio, GP10149-5) for 1 hr at 4°C. The eluates were concentrated using a 30,000 MWCO concentrator (Amicon, UFC503024).

### Affinity-purification of endogenous proteasomal and chaperone-bound complexes

Overnight yeast cultures were inoculated at OD_600_ = 0.25 in 400 mL YPD unless specified otherwise. Cultures were grown to OD600 = 2 and were harvested by centrifugation at 3000 x *g* for 5 min. Cell pellets were washed once with cold water, frozen in liquid nitrogen, and ground with mortar and pestle in the presence of liquid nitrogen. The ground cryo-powders were hydrated for 10 min on ice using proteasome buffer (50 mM Tris-HCl [pH 7.5], 5 mM MgCl_2_, 150 mM NaCl, 1 mM EDTA, 10% glycerol, and protease inhibitors). ATP was added to 1 mM in all buffers used throughout the experiments. Hydrated lysates were centrifuged at 20,000 x *g* for 30 min at 4°C. Affinity-purification using a 3xFLAG affinity tag to each specific bait was conducted by incubating the cleared lysates with 30 μL of anti-FLAG M2-agarose beads (Sigma, A2220-5ML) for 2 hrs at 4°C. The beads were washed twice with 400 μL of proteasome buffer containing 150 mM NaCl with 1 mM ATP, and eluted in 3 beads volumes of proteasome buffer containing 0.2 mg/mL FLAG peptides (GLP Bio, GP10149-5) for 1 hr at 4°C. Affinity-purification using TeV-ProA tag to Rpn11 was conducted by incubating the cleared lysates with 30 μL IgG resin (MP Biomedicals, 855961) for 2 hrs at 4°C. The beads were washed the same way as the FLAG affinity-purification as indicated above. Elution was conducted in 3 beads volume of proteasome buffer containing 1 μL of TEV protease (Promega, PRV6101) for 1 hr at 37°C.

### Native-PAGE analysis

Affinity-purified chaperone-bound complexes (the amount specified in each Figure Legend) were loaded onto 5% discontinuous native gels containing a 2.5% stacking portion. Native gels were electrophoresed for 4.5 hrs at 100V in the cold room. To visualize proteasomal complexes containing GFP-tagged Rpt6 or Rpt1, Native gels were imaged using Amersham Typhoon scanner (GE Healthcare Bio-Sciences AB), using Cy2 filter at a pixel size, 100 μm.

For experiments in Fig. 5C, whole cell lysates (50 μg) were loaded onto 3.5% continuous native gels and electrophoresed for 3 hrs at 100V in the cold room. In-gel peptidase assays were conducted using the fluorogenic peptide substrate LLVY-AMC (Bachem, I-1395.0100) as described previously (32). Native gels were photographed under UV light Bio-Rad Gel Doc EZ Imager to detect AMC fluorescence.

### Heat stress of yeast cell cultures

Overnight yeast cultures were diluted to OD_600_ = 0.25 in synthetic media lacking leucine (marker for plasmids used in Fig. 5C), and were grown to approximately OD_600_ = 1 at 30°C. These cultures were diluted again to OD_600_ = 0.3 in fresh synthetic media lacking leucine in a total volume of 100 mL and further incubated at 37°C for 3.5 hrs to allow for at least two doublings, as in previous studies (30,34,36). Cells were harvested as described in the “Affinity-purification of endogenous proteasomal and chaperone-bound complexes” section, except that whole cell extracts were obtained by centrifuging the hydrated cryo-lysates at 15,000 x g twice for 15 min each in the cold room.

### Immunoblotting

SDS-PAGE or native-PAGE gels were transferred to PVDF membranes (MilliporeSigma, Immobilon, IPVH00010). The PVDF membrane was incubated in 20 mL of blocking buffer, (TBST; Tris-buffered saline containing 0.1% Tween-20), which was supplemented with 5% non-fat dry milk, for 1 hr at room temperature. The membrane was washed twice for 10 minutes using TBST. The membrane was incubated with primary antibodies, which were diluted in blocking buffer. This step was conducted for 1 hr at room temperature for all SDS-PAGE membranes with primary antibodies, and Native-PAGE membranes with anti-Rpt5 antibody. All other primary antibodies for Native-PAGE membranes were incubated overnight at 4°C. Two washes were conducted using TBST as above. The HRP-conjugated secondary antibody at 1:3000 dilutions in blocking buffer (Cytiva, NA931 for anti-Mouse IgG HRP-linked antibody, and Cytiva, NA934 for anti-Rabbit IgG HRP-linked antibody) was incubated with the membranes for 1 hr at room temperature, followed by two washes using TBST. Membranes were then subjected to Enhanced chemiluminescence (Perkin Elmer, Western Blot Chemiluminescence Reagents Plus, NEL105001EA) and were imaged using Bio-Rad ChemiDoc MP Imager.

## DATA AVAILABILITY

All data are contained within the manuscript.

## SUPPORTING INFORMATION

This article contains supporting information (46–48).

## ACKNOWLEDGMENTS

We thank Robb Tomko, Andreas Martin, and Jeroen Roelofs for sharing many plasmids and strains, and James Orth for careful reading of this paper.

## AUTHOR CONTRIBUTIONS

S.S. conducted experiments. S.P. designed the experiments, interpreted results and wrote the manuscript.

## FUNDING AND ADDITIONAL INFORMATION

This study was supported by NIH R01GM127688 to S.P. The content is solely the responsibility of the authors and does not necessarily represent the official views of the National Institutes of Health.

