## Supplemental Figure and Table for "The Penultimate Step of Proteasomal ATPase Assembly is Mediated by a Nas2-Dependent Switch"

Supporting Information contains 2 figures and 5 tables.

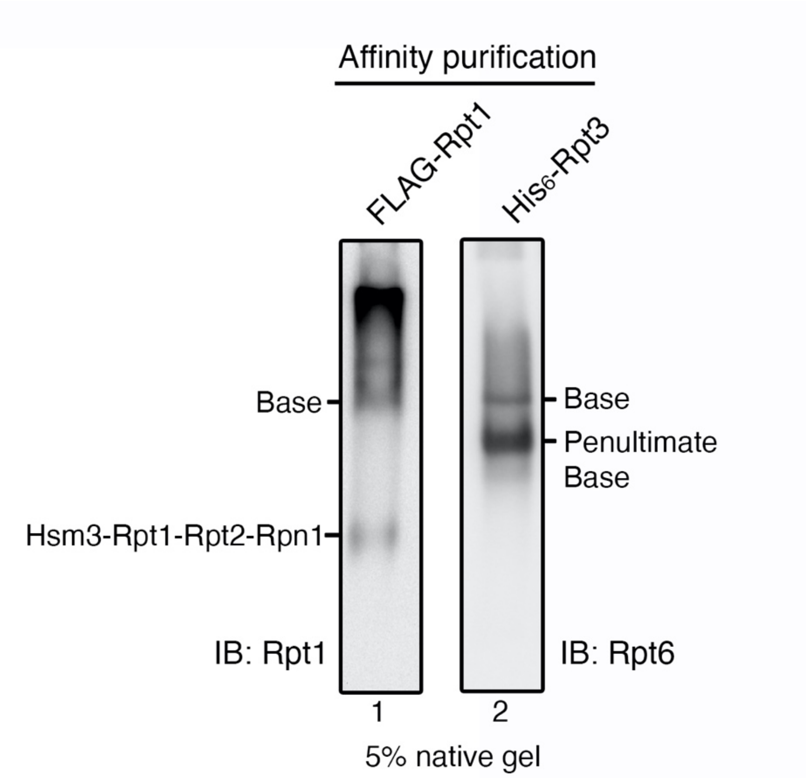


**Figure S1. Additional characterization of the FLAG-Rpt1 and His$_6$-Rpt3 affinity-purification from the heterologous *E. coli* system.**

Affinity-purification was conducted using FLAG-Rpt1 or His$_6$-Rpt3 as a bait, as described in Fig. 1. Upon native gel analysis of FLAG-Rpt1 affinity-purified complexes, the base complex was detected (lane 1; also see Fig. 1, lane 1), together with an additional Rpt1-containing complex, which is likely to be the preceding Hsm3-Rpt1-Rpt2-Rpn1 module, as seen from immunoblotting for its subunit, Rpt1 (lane 1). Upon native gel analysis of His$_6$-Rpt3 affinity-purified complexes, the penultimate base complex was detected (lane 2; also see Fig. 1, lane 2), but the preceding Rpn14-Rpt6-Rpt3-Nas6 module was not detected when examined by immunoblotting for its component, Rpt6, likely due to its rapid progression into the penultimate base complex.

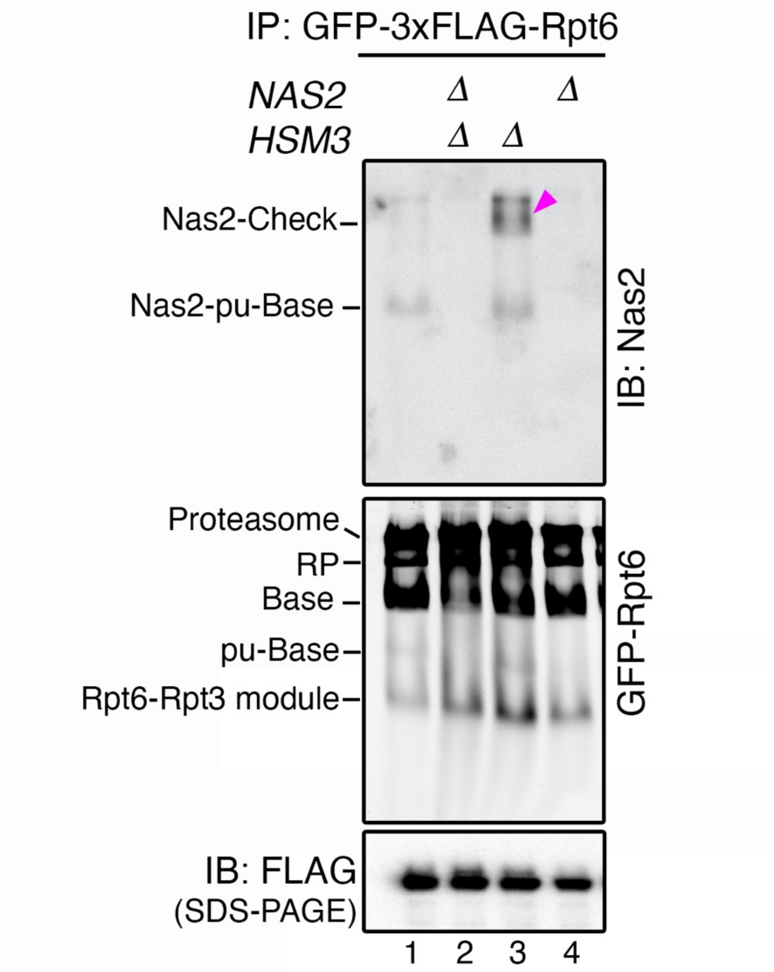


**Figure S2. Validation that the Nas2-Check complex requires the Nas2 chaperone.**

The Nas2-Check complex in the *hsm3Δ* cells (lane 3, arrowhead) forms in a Nas2-dependent manner, since it is no longer detected in the *nas2Δhsm3Δ* double mutants (lane 2). Affinity-purification was conducted using a 3xFLAG affinity tag appended to Rpt6. The purified proteins (7.5 µg) were then subjected to 5% native-PAGE and immunoblotting for Nas2, and GFP fluorescence detection to visualize all Rpt6-containing complexes. Anti-FLAG immunoblot, loading control for FLAG affinity purification.

**Table S1. Base and penultimate base isolated from *E. coli* heterologous system in Fig. 1.**

| | | Subunit Name | No. of unique peptide | No. of total peptide |
|---|---|---|---|---|
| Base complex | Base subunits | *RPN1* | 47 | 336 |
| | | *RPN2* | 56 | 388 |
| | | *RPN13* | 6 | 74 |
| | | *RPT1* | 30 | 382 |
| | | *RPT2* | 29 | 175 |
| | | *RPT3* | 22 | 223 |
| | | *RPT4* | 28 | 203 |
| | | *RPT5* | 30 | 115 |
| | | *RPT6* | 25 | 262 |
| | Chaperones | *NAS6* | 11 | 59 |
| | | *RPN14* | 20 | 194 |
| | | *HSM3* | 24 | 183 |
| Penultimate base complex | Base subunits | *RPN2* | 23 | 26 |
| | | *RPN13* | 4 | 7 |
| | | *RPT3* | 30 | 348 |
| | | *RPT4* | 34 | 285 |
| | | *RPT5* | 33 | 273 |
| | | *RPT6* | 27 | 418 |
| | Chaperones | *NAS6* | 15 | 87 |
| | | *RPN14* | 21 | 364 |
| | | *NAS2* | 13 | 109 |

**Table S2. Endogenous penultimate base and Rpt6-Rpt3 module in Fig. 3B**

| | | Subunit Name | No. of unique peptide | No. of total peptide |
|---|---|---|---|---|
| Endogenous penultimate base complex | Base subunits | *RPN2* | 5 | 5 |
| | | *RPT3* | 4 | 5 |
| | | *RPT4* | 6 | 6 |
| | | *RPT5* | 8 | 8 |
| | | *RPT6* | 9 | 9 |
| | Chaperones | *NAS6* | 3 | 3 |
| | | *RPN14* | 8 | 9 |
| | | *NAS2* | 4 | 4 |
| Endogenous Rpt6-Rpt3 module | Base subunits | *RPT3* | 6 | 9 |
| | | *RPT6* | 7 | 7 |
| | Chaperones | *NAS6* | 3 | 3 |
| | | *RPN14* | 7 | 8 |

### Table S3. Yeast strains used in this study

| Strain | Genotype | Source |
|---|---|---|
| SUB62[a] | *MATa lys2-801 leu2-3, 2-112 ura3-52 his3-Δ200 trp1-1* | (1) |
| SP2017 | *MATa rpt6::Prpt6-yEGFP1F-RPT6-LEU2* | (2) |
| SP4210A | *MATa rpt6::Prpt6-yEGFP1F-RPT6-LEU2. rpt5::rpt5-Δ3 (kanMX6)* | This study |
| SP3027A | *MATα rpt6::Prpt6-yEGFP1F-RPT6-LEU2. nas2::kanMX6* | This study |
| SP4316A | *MATa rpt6::Prpt6-yEGFP1F-RPT6-LEU2 rpn14::hphMX4. hsm3::kanMX6 nas6::HIS3* | This study |
| SP2963A | *MATα rpt6::Prpt6-yEGFP1F-RPT6-LEU2 rpn14::hphMX4. nas6::HIS3* | (2) |
| SP4317 | *MATa rpt6::Prpt6-yEGFP1F-RPT6-LEU2 hsm3::kanMX6 nas6::HIS3* | This study |
| SP4268A | *MATa rpt6::Prpt6-yEGFP1F-RPT6-LEU2 rpn14::hphMX4. hsm3::kanMX6* | This study |
| SP2964B | *MATα rpt6::Prpt6-yEGFP1F-RPT6-LEU2 rpn14::hphMX4* | (2) |
| SP3026A | *MATα rpt6::Prpt6-yEGFP1F-RPT6-LEU2 hsm3::KAN* | (2) |
| SP2965C | *MATa rpt6::Prpt6-yEGFP1F-RPT6-LEU2 nas6::his3* | (2) |
| SP3025A | *MATα rpt6::Prpt6-yEGFP1F-RPT6-LEU2 nas2::KAN hsm3::KAN* | (2) |
| SP2781A | *MATα rpt1::*$P_{rpt1}$*-yEGFP1F-RPT1-LEU2* | (2) |
| SP3033A | *MATa rpt1::*$P_{rpt1}$*-yEGFP1F-RPT1-LEU2 nas2::KAN* | (2) |
| SP3032A | *MATa rpt1::*$P_{rpt1}$*-yEGFP1F-RPT1-LEU2 hsm3::KAN* | (2) |
| SP3031A | *MATα rpt1::*$P_{rpt1}$*-yEGFP1F-RPT1-LEU2 nas2::KAN hsm3::KAN* | (2) |
| SP1655A | *MATa nas2::NAS2-6×Gly-3×FLAG (kanMX6)* | (3) |
| SP3210 | *MATα nas2::NAS2-6×Gly-3×FLAG(kanMX6) nas6::HIS3 rpn14::hphMX hsm3::KAN* | This study |
| SP3198A | *MATa nas2::NAS2-6×Gly-3×FLAG (kanMX6) nas6::HIS3 rpn14::hphMX* | This study |
| SP3196 | *MATa nas2::NAS2-6×Gly-3×FLAG (kanMX6) nas6::HIS3 hsm3::KAN* | This study |
| SP3194 | *MATα nas2::NAS2-6×Gly-3×FLAG (kanMX6) rpn14::hphMX hsm3::KAN* | This study |
| SP3200A | *MATa nas2::NAS2-6×Gly-3×FLAG (kanMX6) rpn14::hphMX* | This study |
| SP3202 | *MATα nas2::NAS2-6×Gly-3×FLAG (kanMX6) nas6::HIS3* | This study |
| SP3193 | *MATα nas2::NAS2-6×Gly-3×FLAG (kanMX6) hsm3::KAN* | This study |
| SP1677A | *MATa nas6::NAS6-6× Gly-3×FLAG (hphMX)* | (3) |
| SP3128A | *MATα nas6::NAS6-6×Gly-3×FLAG (hphMX) rpn14::hphMX hsm3::KAN nas2::NAT* | (4) |
| SP2692A | *MATa nas6::NAS6-6×Gly-3×FLAG (hphMX) rpn14::hphMX hsm3::KAN* | (4) |
| SP3124A | *MATa nas6::NAS6-6×Gly-3×FLAG (hphMX) rpn14::hphMX nas2::NAT* | (4) |
| SP3127A | *MATa nas6::NAS6-6×Gly-3×FLAG (hphMX) hsm3::KAN nas2::NAT* | (4) |
| SP3129A | *MATα nas6::NAS6-6×Gly-3×FLAG (hphMX) hsm3::KAN* | (4) |
| SP1883A | *MATa nas6::NAS6-6×Gly-3×FLAG (hphMX) rpn14::hphMX* | (4) |
| SP3132A | *MATα nas6::NAS6-6×Gly-3×FLAG (hphMX) nas2::NAT* | (4) |
| SP1493A | *MATa rpn2::RPN2-3×FLAG:HIS3* | This study |
| SP4458A | *MATa rpn2::RPN2-3×FLAG:HIS3 rpn14::hphMX hsm3::KAN* | This study |
| SP4457A | *MATa rpn2::RPN2-3×FLAG:HIS3. hsm3::KAN* | This study |
| SP4456B | *MATa rpn2::RPN2-3×FLAG:HIS3. rpn14::hphMX* | This study |
| sDL133 | *MATa rpn11::RPN11-TEV-ProA (HIS3)* | (5) |
| SP3111A | *MATα rpn11::RPN11-TEV-ProA (HIS3) hsm3::KAN nas2::NAT* | This study |
| SP3109A | *MATa rpn11::RPN11-TEV-ProA (HIS3) hsm3::KAN* | This study |
| SP3107A | *MATa rpn11::RPN11-TEV-ProA (HIS3) nas2::NAT* | This study |
| SP4245A | *MATa nas2::NAS2-6×Gly-3×FLAG (kanMX6) rpt6::HIS3. [YCplac33-RPT6]* | This study |
| SP4246A | *MATa nas2::NAS2-6×Gly-3×FLAG (kanMX6) rpt3::HIS3. [YCplac33-RPT3]* | This study |
| SP1654B | *MATa nas2::KAN* | (4) |

[a]All strains are isogenic to SUB62 genetic background.

### Table S4. Plasmids used in this study

| Name | Description | Source and Reference |
|---|---|---|
| N/A | FLAG-Rpt1, Rpt2, $His_6$-Rpt3, Rpt4, Rpt5, Rpt6 in pCOLA-1 | Martin Laboratory (6) |
| N/A | Rpn14, Nas6, Nas2, Hsm3 and tRNAs for rare codons in pACYCDuet-1 | Martin Laboratory (6) |
| N/A | Rpn1, Rpn2, Rpn13 in pETDuet-1 | Martin Laboratory (6) |
| N/A | FLAG-Rpt1, Rpt2, $His_6$-Rpt3, Rpt4, Rpt5 without last 5 amino acids, Rpt6 in pCOLA-1 | Martin Laboratory (6) |
| pSP310 | Rpn14, Nas6, Nas2 with a premature stop codon, Hsm3 and tRNAs for rare codons in pACYCDuet-1 | This study |
| pSP151 | $His_6$-Rpt5 in pRSF-Duet-1 | (4) |
| pSP158 | Untagged Rpt4 in pRSF-Duet-1 | (4) |
| pSP128 | GST-Nas2 in pGEX6P-1 | (4) |
| pJR751 | pRS316-*RPT1* | Roelofs Laboratory (2) |
| pRT357 | pRS314-*RPT1* | Tomko Laboratory (7) |
| pRT702 | YCplac111-*RPT2* | Tomko Laboratory (8) |
| pRT364 | YCplac111-*RPT3* | Tomko Laboratory (8) |
| pRT1528 | YCplac111-*RPT4* | Tomko Laboratory (7) |
| pRT1408 | pRS315-*RPT5* | Tomko Laboratory (7) |
| pRT1496 | YCplac111-*RPT6* | Tomko Laboratory (7) |
| pRT1409 | pRS314-*rpt1(E310Q)* | Tomko Laboratory (7) |
| pRT1410 | YCplac111-*rpt2(E283Q)* | Tomko Laboratory (7) |
| pRT1411 | YCplac111-*rpt3(E273Q)* | Tomko Laboratory (7) |
| pRT1529 | YCplac111-*rpt4(E283Q)* | Tomko Laboratory (7) |
| pRT1413 | pRS315-*rpt5(E282Q)* | Tomko Laboratory (7) |
| pRT1497 | YCplac111-*rpt6(E249Q)* | Tomko Laboratory (7) |

### Table S5. Antibodies used in this study

| Name | Source and reference |
|---|---|
| Rabbit polyclonal Anti-Nas2 | Roelofs Laboratory (9) |
| Rabbit polyclonal Anti-Rpt5 | Enzo Lifesciences, BML-PW8245 |
| Rabbit polyclonal Anti-Rpt3 | Enzo Life Sciences, BML-PW8250 |
| Rabbit polyclonal Anti-Rpt6 | Carl Mann laboratory (10) |
| Rabbit polyclonal Anti-Rpt1 | Carl Mann laboratory (10) |
| Rabbit polyclonal Anti-Rpn14 | Finley laboratory (11) |
| Mouse monoclonal Anti-FLAG | Sigma, F3165 |
| Mouse monoclonal Anti-His | Sigma, H1029 |
| Mouse monoclonal Anti-Pgk1 | Life Technologies, 459250 |
